# *aer2vec*: Distributed Representations of Adverse Event Reporting System Data as a Means to Identify Drug/Side-Effect Associations

**DOI:** 10.1101/780544

**Authors:** Jake Portanova, Nathan Murray, Justin Mower, Devika Subramanian, Trevor Cohen

## Abstract

Adverse event report (AER) data are a key source of signal for post marketing drug surveillance. The standard methodology to analyze AER data applies disproportionality metrics, which estimate the strength of drug/side-effect associations from discrete counts of their occurrence at report level. However, in other domains, improvements in predictive modeling accuracy have been obtained through representation learning, where discrete features are replaced by distributed representations learned from unlabeled data. This paper describes aer2vec, a novel representational approach for AER data in which concept embeddings emerge from neural networks trained to predict drug/side-effect co-occurrence. Trained models are evaluated for their utility in identifying drug/side-effect relationships, with improvements over disproportionality metrics in most cases. In addition, we evaluate the utility of an otherwise-untapped resource in the Food and Drug Administration (FDA) AER system – reporter designations of suspected causality – and find that incorporating this information enhances performance of all models evaluated.

## Introduction

The need for post-marketing surveillance of the unintended effects of pharmaceutical products has been repeatedly highlighted by the withdrawal of drugs on account of previously undetected serious side effects^1^. Notoriously, the widely utilized Cox-2 inhibitor Vioxx (rofecoxib) was removed from market five years after release as it was shown to significantly increase the risk of myocardial infarction, and was estimated to have caused between 88,000 and 140,000 coronary cardiac events in the United States alone while on market^2,3^. More recently proton pump inhibitors, many of which are available without prescription, have been shown to significantly increase risk of bone fracture as well as chronic kidney disease^4,5^. These findings are not atypical – between the years of 2001 and 2010, nearly one third of drugs approved by the US Food and Drug Administration (FDA) had a subsequent safety event in the form of a label change, safety communication or withdrawal^6^. The median time from drug release to detection of such events was 4.2 years^6^. The morbidity and mortality caused by previously undetected drug side effects could be mitigated by earlier detection, as could the societal costs of such adverse events – estimated at 3.5 billion dollars in 2006^7^. One solution to the inherent shortcomings of clinical trials in detecting adverse drug events (ADEs) is improving their identification after release to market. Consequently, post-marketing surveillance through pharmacovigilance, defined as “the study of the safety of marketed drugs under the practical conditions of clinical use in large communities”, is an essential component of drug safety^8^.

Traditionally, such safety surveillance has involved the analysis of reports of suspected ADEs submitted by healthcare practitioners, pharmaceutical companies and patients. In the United States the FDA maintains the FDA Adverse Event Reporting System (FAERS), providing a database of ADE reports from as early as 1969^2^. Adverse events in the AERS are reported by healthcare professionals, consumers, and pharmaceutical companies. Each report includes one or more adverse events that appear to be associated with the administration of a drug as well as other drugs prescribed to the patient concerned and their therapeutic indications. Of importance to the current research, reporters have the opportunity to indicate which of a set of prescribed drugs they suspect caused the ADE under consideration, by designating these drugs as primary (presumed cause) or secondary (potential cause) suspects. Large numbers of reports exist in this repository, with over a million received in the year 2014 alone^3^. Consequently, automated methods of analysis are a prerequisite to the identification of actionable safety signals.

An important component of post-marketing drug surveillance is the identification of statistically significant drug/side-effect association, termed “signal detection”, and considerable research has been devoted to the development and evaluation of methods for this purpose^4,5,6,7,8,9^. In order to identify meaningful associations from large and unlabeled data sets such as AERS reports, data mining techniques known as Signal Detection Algorithms (SDAs) are employed^10^. SDAs can be subdivided into two main classes, *disproportionality analysis* (DPA) and *multivariate modeling*^10^. DPA methods quantify the extent to which drugs and side-effects are reported together beyond what would be expected by chance. For a review of DPA methodologies, we refer the interested reader to Bate et al^9^. Multivariate modeling can mitigate for confounding polypharmacy variables as well as the lack of a quantitative drug prescription frequency. SDAs are the most frequently utilized methods used in analyzing AERS data, and they are well documented in the post-marketing drug surveillance literature.^6,7,8,9,10^

While SDAs have shown their utility as a means to identify safety signals from adverse event reports, they are not without limitations. In particular, they lack the capacity to draw connections between similar drugs (e.g. the entire family of selective cox-2 inhibitors) and related side-effects (e.g. myocardial infarction and other cardiac events such as stroke) as a means to enhance the strength of safety signals for relatively rare events. This situation is analogous to recent developments in natural language processing (NLP), where discrete ‘one-hot’ vector representations of words have been largely superseded by distributed vector representations, which are learned from a large unlabeled corpus such that words that occur in similar contexts have similar vectors. In the current paper we adapt skip-gram neural embeddings, a widely used representation learning technique in NLP embodied in the popular *word2vec* software package, to the task of representing drugs and side-effects appearing in FAERS data and evaluate their utility as a means to detect safety signals.

Specifically, this paper describes *aer2vec*, a novel representational approach for adverse event report data in which a neural network is trained to predict drug/side-effect co-occurrence events. The trained model is evaluated for its utility as a means to identify causal drug/side-effect relationships. Our approach leverages methods of distributional semantics to represent the AERS database as a vector space. Distributional semantics methods - such as the neural embeddings implemented by word2vec - attempt to model the semantic similarity and relatedness between words. These methods are based on the distributional hypothesis which states that words that occur in similar contexts tend to have similar meanings^11^. A broad range of methodological approaches have been applied to learn word representations from text (for reviews see ^12,13^). A recent trend involves the application of neural-probabilistic models, such as the skipgram and continuous-bag-of-words architectures embodied in the popular *word2vec* and *fastText* software packages^14,15,16,17^. As implied by the term “neural-probabilistic”, these models are trained to predict the occurrence of a context word given an observed term. Although generally utilized during training only, this probabilistic aspect of the model can be used to recover learned probabilities for observing one word in the context of another. It is this aspect of neural-probabilistic models that we adapt to represent AERS data in the current work.

Our primary hypothesis in conducting this work was that, at least in some cases, the capacity of distributional semantics models to generalize between similar drugs and side-effects may lead to improved performance in the task of identifying drug/side-effect relationships. A secondary hypothesis was that restricting the data considered by our models, and perhaps baseline disproportionality metrics also, to designations of primary and/or secondary suspect may improve their performance.

## Methods

### Disproportionality metrics

We compare the performance of our models to two widely-used disproportionality metrics, the Proportional Reporting Ratio (PRR) and the Reporting Odds Ratio (ROR) ^4,18^. Disproportionality metrics are derived from 2×2 contingency tables (Table 1) constructed using report level statistics:

**Table 1.** 2×2 Contingency Table. The cells indicate counts of co-occurrence events in reports.

|  | ADE of Interest | Other ADE's |
| --- | --- | --- |
| Drug of Interest | a | b |
| Other Drugs | c | d |

From this table, we derive two disproportionality metrics, the PRR and the ROR. The PRR estimates the probability of an ADE given a drug divided by the probability of this ADE without the drug, or P(ADE | ~drug) / P(ADE | ~drug). This probability can be calculated from the 2×2 table as follows:

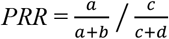

In statistical terms, the ROR is the ratio of odds of an ADE occurring vs not occurring given a drug to the odds of this ADE occurring vs not occurring given other drugs. It is calculated from the 2×2 table as follows:

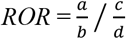

These disproportionality metrics provide a baseline for our evaluation.

### Skipgram-with-negative-sampling

The skipgram-with-negative-sampling (SGNS) algorithm of Mikolov and his colleagues trains a shallow neural network to predict the probability of a context word *c* occurring nearby to an observed word *w*, P(*c*|*w*)^19^. For example, given the sliding window “adverse [drug] events”, the model would be trained to optimize P(*adverse*|*drug*) and P(*events*|*drug*). While it would in theory be possible to train this model using a softmax objective in which all other unique terms in the corpus are considered as counterexamples to the observed context term in every window, this would be computationally intractable. The SGNS algorithm provides a practical way to train neural embeddings by instead considering as counterexamples a small number (usually 5-15) of *negative samples*, randomly drawn words that probably do not occur with the focus word within a sliding window. More formally, the optimization objective of the SGNS algorithm can be expressed as follows:^20^

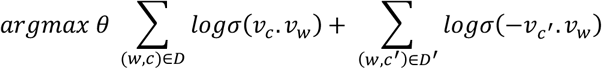

where *w* are observed words in a set of documents *D*, and *c* and *c’* are context terms that occur with these observed terms in a sliding window or are randomly drawn as counterexamples respectively. v_w_ and v_c_ are vector representations of these terms, specifically the input weights (v_w_) and output weights (V_c_) of the neural network for each term in the vocabulary. σ is the sigmoid function, which converts the scalar product between these vectors into a value between one and zero which can be interpreted probabilistically. Training in SGNS occurs through stochastic gradient descent with a linearly decreasing learning rate. The algorithm has several hyper-parameters that have been shown to influence performance across tasks^21^. Of importance for the current paper, the number of negative sample terms drawn as counterexamples to each observed term is a parameter of the model, and these terms are drawn with a probability derived from the frequency, *f*, with which they appear in the corpus (i.e. *f* = *count* / *total number of non-unique terms*), *f ^.75^*. Another hyper-parameter setting concerns *subsampling* – ignoring terms occurring above a predetermined frequency threshold *t* with probability 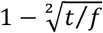. With SGNS, the input weights are usually retained as word embeddings and used in downstream tasks, and the output weights are discarded. However, P(*c*|*w*) can be estimated by retaining these weights and calculating σ(*v*_*c*_. *v*_*w*_).

### aer2vec

With *aer2vec*, we adapt the SGNS algorithm to estimate probabilities for drug/side-effect relations from AERS data. We derive two neural architectures, *aer2vec+* which estimates P(drug|ADE) and *aer2vec−* which estimates P(ADE|drug). Our motivation in developing distinct architectures for each of these estimates was to ensure that only information pertinent to an estimate of interest is encoded by the architecture responsible for it, eliminating one channel through which noise might be introduced. In addition we wished to determine which of these two possible neural-probabilistic estimates is of greater utility for identification of drug/ADE relationships.

These architectures are illustrated in Figure 1, which shows a simplified *aer2vec* architectures for 8 side effects and 10 drugs. As is the case with the original skipgram architecture, each observed term (in this case a drug) (aer2vec+) or ADE (aer2vec−) is connected to a hidden layer by input weights, which are in turn connected to a predicted term by output weights. While the output weights are usually discarded when generating word embeddings, we retain them to facilitate predicting P(drug|ADE) and P(ADE|drug) in *aer2vec+* and *aer2vec−* respectively. When trained with negative sampling, the aer2vec architectures have the following optimization objectives:

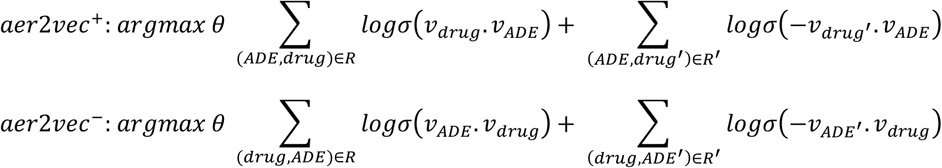

**Figure 1:**
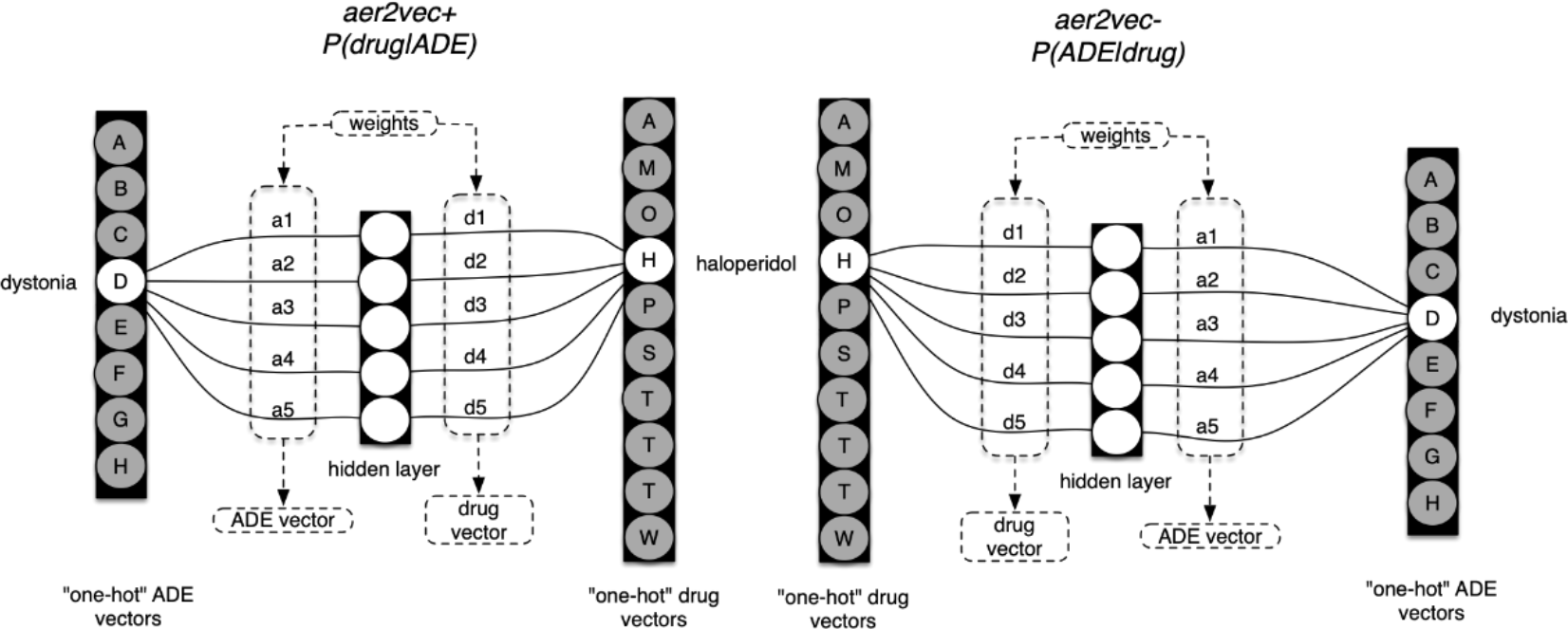
Two mirror-image aer2vec architectures, *aer2vec+* (left) and *aer2vec−* (right).

This is similar to SGNS with word windows, with the exceptions that (1) co-occurrence occurs at the level of a report (R) rather than a sliding window; and (2) drugs and ADEs have either input weight vectors or output weight vectors depending on the model, whereas words in neural word embeddings have both input and output weights. With aer2vec, both sets of weights are retained, permitting estimation of both P(drug|ADE) with aer2vec+, and P(ADE|drug) with aer2vec−, as σ(*v*_*drug*_.*v*_*ADE*_).

## Evaluation

### AER data

As data for all models, we used a standardized version of the FAERS data set released by Banda et al^22^. This data set contains standardized versions of the FAERS data for the years 2004 to 2013. Standardization here indicates that drugs are mapped to RxNorm concepts, and the side-effects to SNOMED-CT^22^. We extracted three component data sets. First, we generated an unconstrained data set (FULL), including all the drugs that were mapped to RxNorm and side effects that occur in the “pt” field of any report with them. We then use the role code variable to produce two subsets of this data set. The role codes we used were primary suspect (PS) and secondary suspect (SS). These terms denote whether a drug is the primary or secondary suspect for a side-effect, and we leveraged these to produce data subsets consisting of only drugs and side-effects in primary suspect relationships (PS), and the disjunction of primary and secondary suspect relationships (PS-SS).

### aer2vec models

For each of the three dataset configurations, we generated *aer2vec+* and *aer2vec−* models. All models used 500-dimensional vectors and were trained for 25 iterations across the data with five negative samples per positive example, and without subsampling of frequently occurring concepts. We did not attempt to optimize these hyperparameters in our initial experiments. During training we encoded all of the drug and ADE terms meeting the relevant constraints (PS/PS-SS/FULL), aside from a stoplist of 28 ADE terms (such as “NA”, “test”, “error”) that were identified as lacking semantic content upon inspection of the data.

### Evaluation sets

A number of research groups have released reference standards that provide a uniform testing ground for pharmacovigilance systems. In some cases, these are manually curated reference sets of positive and negative relationships between drugs and potential ADEs^23,24^. For example, the widely-used OMOP reference standard produced by Ryan and his colleagues consists of 165 positive and 234 negative examples of drug/ADE relationships, spanning four serious side-effects (such as renal failure), and developed through extensive manual review of the literature and other sources^23^. These sets have been extensively curated, so there is reason to believe they are largely accurate (for positive examples in particular – some amendments have been suggested for negative controls). We evaluated aer2vec for its ability to distinguish between positive and negative controls in the reference standards produced by Ryan *et al* (henceforth, OMOP) and Coloma *et al* (henceforth EU-ADR) containing 399 and 94 examples respectively^23,24^. For both disproportionality metrics and *aer2vec* models, a small number of examples were eliminated from the FULL (n=1) and PS/PSSS (n=6) configurations of the OMOP set respectively. These examples concerned the drugs “olmesartan_medoxomil” (all configurations), “endopeptidases” and “alatrofloxacin” (PS/PSSS configurations), which are not present in the source data. All examples from the EU-ADR set (n=94) were retained.

### Quantitative evaluation

For each drug/event pair in the two reference standards (OMOP/EU-ADR) under each of the three configurations (PS/PS-SS/FULL), we calculated disproportionality metrics (PRR/ROR), as well as P(drug|ADE) and P(ADE|drug) with *aer2vec+* and *aer2vec−* respectively. These scores were used to estimate the area under receiver operator characteristic curve (AUROC) and area under the precision recall curve (AUPRC) to provide a basis for comparison. In order to account for variance in the stochastic initialization of the *aer2vec* embedding weights, we retrained each *aer2vec* model 10 times, and report mean area under curve (AUC) across these runs.

### Qualitative evaluation

In order to assess the likely utility of the vector embeddings generated during the course of training our models for downstream supervised machine learning, we include some examples of nearest neighbor searches amongst the input and output embeddings for the best-performing model.

### Hyperparameter settings

Subsequent to our initial experiments, we explored the influence of two *aer2vec+* hyperparameters on model performance. We repeated our initial experiments with the PS and FULL configurations at dimensionality of 100, 250, 500 (the original setting) and 1,000 dimensions; and at 100 dimensions with subsampling thresholds of 10^−3^, 10^−4^, 10^−5^ and without subsampling (the original setting). For these experiments we used 5 (rather than the original 25) epochs of training.

## Results

### Quantitative evaluation

Disproportionality metric results for all three versions of the data sets across both reference standards, are shown in Table 2. These results provide points of comparison for *aer2vec*. As anticipated (see for example Waller et al ^25^), these metrics exhibit similar performance. Of note, both baseline metrics show an increase in performance when reducing the data set by role code. In the OMOP set, restricting to the PS-SS leads to best performance with AUROC’s of.744 for both PRR and ROR. In the EU-ADR set, best performance is obtained with the PS constraint with AUROC’s of .935 for both methods. Across both reference sets and disproportionality metrics, there is an absolute AUROC increase of approximately 10% when we reduce the full data set to the primary suspects.

**Table 2:** Performance of disproportionality metrics. PS-SS= primary and secondary suspects. PS=primary suspects. Best results for each metric on each reference set are in boldface.

|  |  | OMOP |  |  | EU-ADR |  |  |
| --- | --- | --- | --- | --- | --- | --- | --- |
|  |  | FULL | PS-SS | PS | FULL | PS-SS | PS |
| PRR | AUROC | 0.646 | **0.744** | 0.742 | 0.841 | 0.918 | **0.935** |
|  | AUPRC | 0.580 | **0.694** | 0.729 | 0.836 | 0.931 | **0.941** |
| ROR | AUROC | 0.646 | **0.744** | 0.742 | 0.843 | 0.919 | **0.935** |
|  | AUPRC | 0.579 | **0.691** | 0.723 | 0.840 | 0.932 | **0.941** |

### aer2vec

Results for the two aer2vec models are shown in Table 3. These results show improvement over disproportionality metrics for both reference sets in every configuration (FULL,PS-SS and PS) by both metrics of evaluation. As was the case with the disproportionality metrics, the results show an improvement in AUC when reducing the data set by role code, with an absolute AUC increase of approximately 10% when we constrain the FAERS data to primary suspects. While *aer2vec+* and *aer2vec−* perform similarly to one another on the EU-ADR set, the *aer2vec+* model performs considerably better on the larger OMOP set. As was the case with disproportionality metrics, performance improves by around 10% (absolute AUROC) when restricting data to primary suspect relationships only. In addition, the best *aer2vec* results exceed the best disproportionality metric result on both data sets, with substantive improvements in performance across all configurations with the OMOP set in particular. As indicated by the confidence intervals of these mean values, performance was remarkably consistent across iterations. Across these ten iterations, all differences in performance between PS, PS-SS and FULL configurations of the same model were statistically significant by an unpaired t-test, as were all differences between *aer2vec+* and *aer2vec−* models in the same configuration, aside from the sole case of the PS-SS configuration with the EU-ADR set.

**Table 3:**
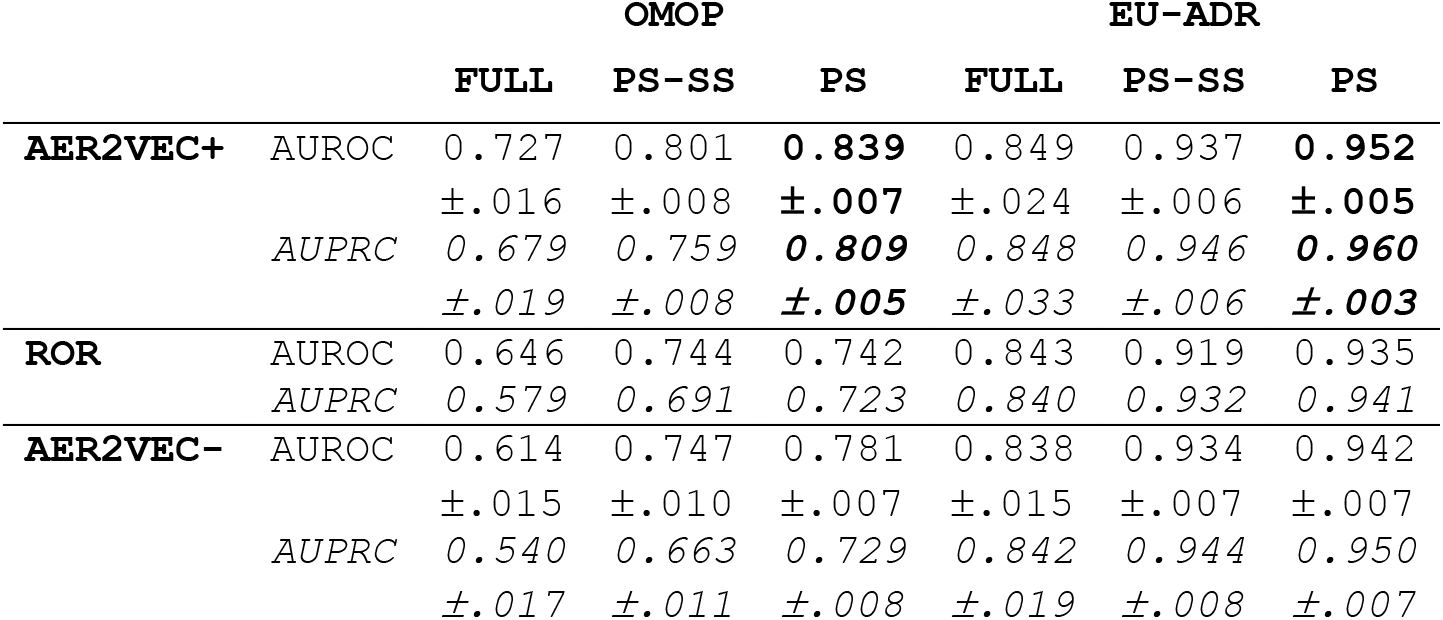
*aer2vec* results. Mean AUROC/AUPRC ±1.96*STD across 10 repeated runs. PS-SS= primary and secondary suspects. PS=primary suspects. Best results for each metric on each reference set are in boldface. The best baseline model (ROR) is shown in the middle two rows for the purpose of comparison.

Table 4 illustrates three ways in which the trained embedding space (in this case *aer2vec+* PS) can be interrogated. The first two results columns involve comparing the same types of entities – side effects (in this case input weight embeddings) or drugs (in this case output weight embeddings). Furthermore, it is possible to recover predictions for observing a particular drug given a side effect has been reported as shown in the third results column. As indicated in the accompanying comments, the results are interpretable in many cases and suggest the resulting embeddings may be of utility for downstream supervised learning models on account of the similarities they capture.

**Table 4:**
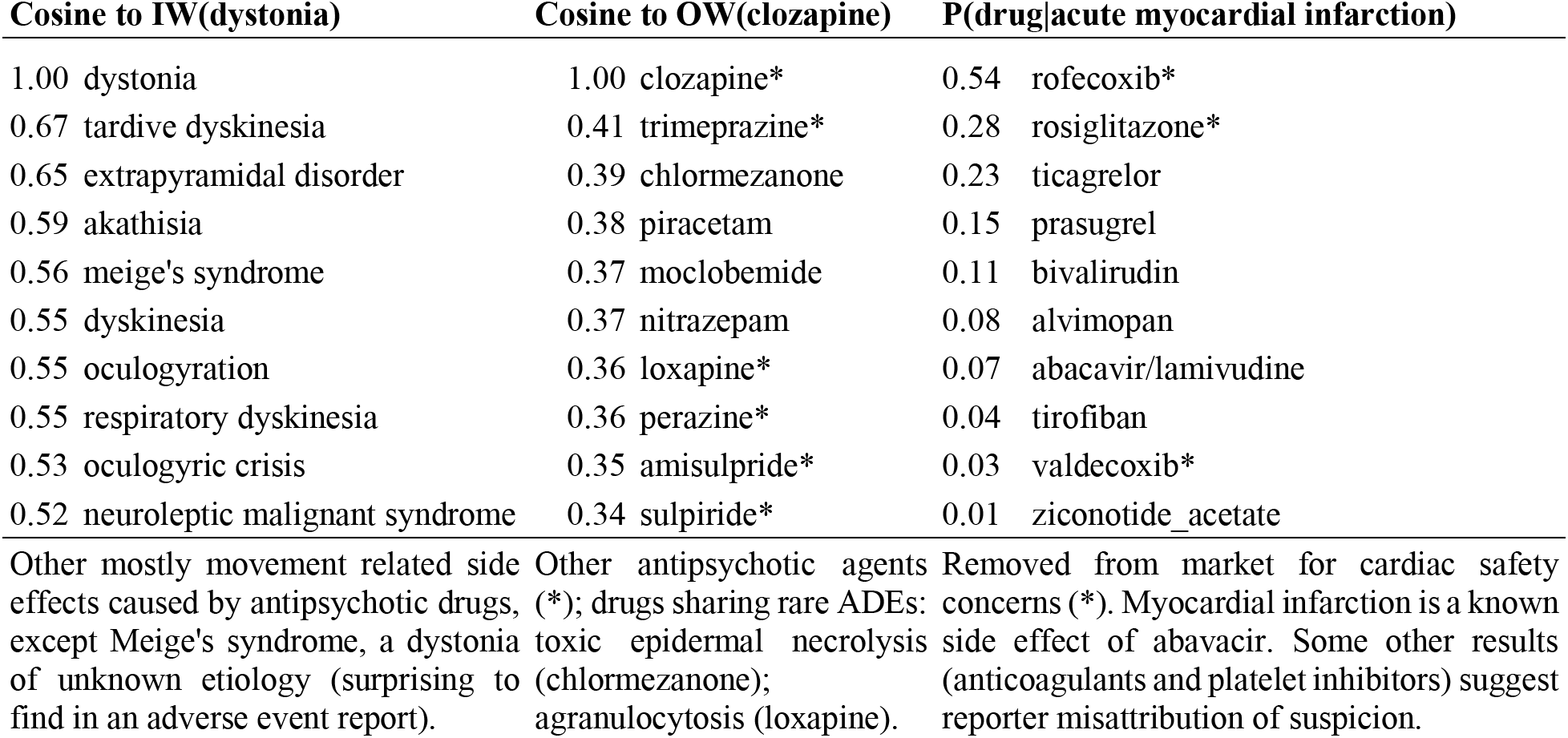
Nearest neighbors in *aer2vec+* PS space. IW=input weights, OW=output weights

These results also give some insight into the possible mechanisms underlying *aer2vec’s* improvements in performance over established disproportionality metrics, as the estimation of P(drug|ADE) (or vice-versa) will be influenced by similarities between drug and side-effect terms. For example, P(drug|tardive dyskinesia) will be elevated for a drug reported with the term “extrapyramidal disorder”, which is a useful generalization in this case because of the hyponymic relationship between these side effects (tardive dyskinesia is a type of extrapyramidal disorder). More broadly, one would anticipate the model inferring that drugs that share some side effects may share others.

### Hyperparameter settings

Evaluation of performance across different hyperparameter settings reveals that subsampling impairs performance (Figure 2, left) and consistency across a range of dimensions (Figure 2, right). Best performance across these settings is shown in Table 5, with improvements in OMOP set performance in particular.

**Figure 2:**
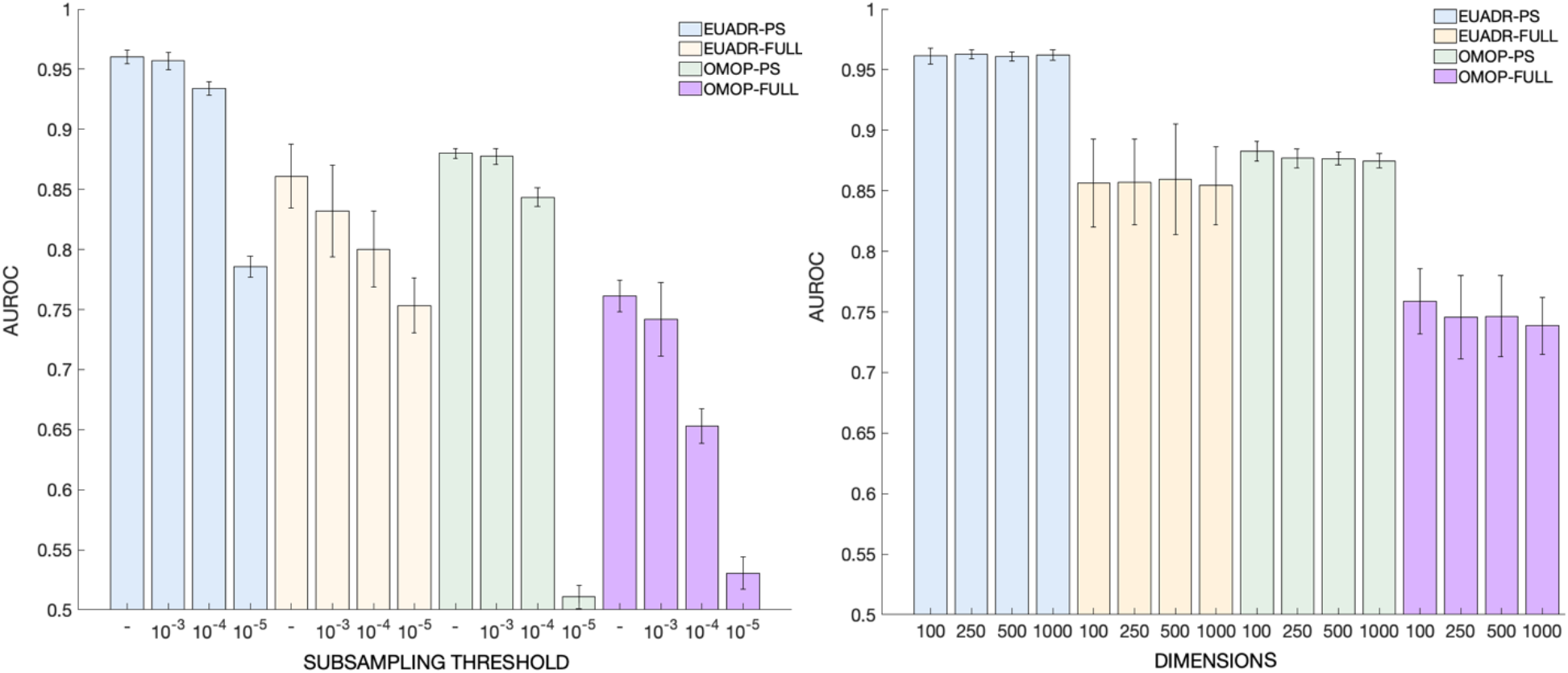
Mean AUROC (n=10) with 95% CI at different subsampling thresholds (left; ‘-’ indicates no subsampling) and dimensionalities (right). 0.5 baseline indicates performance with random ordering.

**Table 5:** Best aer2vec+ performance across hyperparameter settings (100 dimensions, no subsampling)

|  |  | OMOP FULL | OMOP PS | EU-ADR FULL | EU-ADR PS |
| --- | --- | --- | --- | --- | --- |
| **AER2VEC+** | AUROC | 0.761±.013 | **0.880±.004** | 0.861±.026 | **0.960±.006** |
|  | AUPRC | 0.712±.021 | **0.844±.007** | 0.863±.038 | **0.968±.008** |

## Discussion

Our results demonstrate improved signal detection from the FAERS data when applying an adaptation of the SGNS algorithm to represent observational data, providing a novel methodology for the identification of drug/side-effect associations. Across both reference standards and three different configurations of our source dataset, P(drug|ADE) estimates produced by the *aer2vec+* model consistently result in a higher AUC than was obtained by either of the two disproportionality metrics evaluated. The *aer2vec−* model also outperformed disproportionality metrics in most but not all cases. Furthermore, we found that the role code information in the FAERS data can be used to subset these data in a way that improves performance. Lastly, we discovered that *aer2vec+* outperforms *aer2vec−* in all cases. These findings have implications for the analysis of adverse event report data.

Our primary finding is that *aer2vec*+ outperforms the standard disproportionality metrics across two reference sets and all dataset configurations, with improvements in performance of up to 10% in absolute AUROC on the larger and more challenging OMOP set, and smaller but still substantial improvements in performance on the EU-ADR set. This supports our initial hypothesis that distributed representations (concept embeddings) derived from AERS data may be of value for pharmacovigilance. Qualitative analysis of neighboring embeddings suggests that trained embeddings capture regularities in the data. Concepts occurring in similar AER contexts have similar embeddings, permitting our models to generalize across related drugs and ADEs, suggesting a mechanism underlying their enhanced performance. In the current work, we applied these embeddings by recapitulating the probabilistic estimates used during the course of training. However, the benefits of pre-trained distributed representations are often best demonstrated in downstream supervised learning. In our previous work we have shown that literature-derived embeddings used as a basis for supervised machine learning produce state-of-the-art performance (AUROC=.96) on the OMOP reference set^26,27^. A logical next step will involve evaluating the utility of *aer2vec* embeddings as a complement to literature-derived embeddings for supervised machine learning. Intriguingly, best performance with *aer2vec* on the EU-ADR set exceeds that obtained with supervised models - both in our work^26,27^ and when classifiers were trained on manually engineered features derived from a range of sources including the biomedical literature, FAERS data and pharmaceutical product labels^28^. One reason for this may be the limited number of training examples per side-effect available relative to the OMOP set, where recapitulating *aer2vec’s* neural-probabilistic learning objective does not in and of itself lead to state-of-the-art performance. In addition, prior work leveraging side-effect patterns in AERS data for drug repurposing suggests *aer2vec* embeddings that represent such patterns may be leveraged for this purpose also^29^.

A second notable finding is that constraining the data set in accordance with human judgment of suspected causality enhances performance. The role code variable in the FAERS data set provides valuable information that has generally not been used in prior published work, and leveraging this information to constrain the occurrence events in the data resulted in substantive improvements in performance for all models evaluated. It seems likely that the mechanism in this case involves reducing the influence of confounding variables, providing a useful addition to the armament of methods that have been deployed for this purpose in pharmacovigilance^30,31,32^. However, incorporation of human judgment into statistical models introduces a degree of subjectivity that warrants further discussion. One might argue that attempts to leverage these designations of suspicion would accentuate documented biases in spontaneous reporting, such as *notoriety bias* -a tendency toward increased reporting of adverse events that have recently appeared in safety alerts^33^. A more pragmatic perspective might be to consider the cumulative independent judgments of a diverse group of adverse event reporters – who are in some cases domain experts and have access to information concerning temporality and other factors that are not explicit in reporting data alone – as exemplifying the “wisdom of crowds” ^34^. At a minimum, the utility of reporter designations of suspected causality as a means to improve predictive modeling performance across multiple models and two reference standards suggests a need for further investigation into the circumstances in which this information is of value for post-marketing drug surveillance.

In addition, we find that *aer2vec+* performs better than *aer2vec−*. While both approaches outperform the disproportionality metrics in most cases, *aer2vec+* is consistently better than *aer2vec−*, with better mean AUC scores for *aer2vec+* across both reference sets in all configurations. We reason that predicting the drug given the side-effect is the more suitable approach to identifying drug/side-effect associations from AERS data, and that future research should prioritize *aer2vec+* as a means to represent these data for pharmacovigilance purposes. Subsequent exploration of model hyperparameters showed subsampling of frequent terms reduced performance, which seems intuitive given the nature of the task. In addition, these experiments revealed improved *aer2vec+* performance on the OMOP set in particular at a lower dimensionality than in our original experiments. Optimal performance at lower dimensionality has been observed in other distributional semantic models also, and we refer the interested reader to Yin and Shen’s recent theoretical account of the relationship between dimensionality and task performance in such models^35^. Code to reproduce our experiments is publicly available^i^, as are trained models for our primary and best results^ii^.

## Limitations

An important limitation of this study concerns the use of reference sets containing well-established side effects. More stringent evaluations using time-indexed sets are needed to determine performance with emerging side effects^36^. These models were also created with standardized data. Our methods have yet to be evaluated in the context of raw FAERS data, although one might hypothesize that the capacity for generalization provided by distributed representations would offer greater advantages in the context of data that have not been normalized. Another limitation is that this data set only contains data up to 2015. We plan to address these limitations by training our models using raw FAERS data in future experiments. Finally, we compared *aer2vec* performance to two disproportionality metrics only. Outperforming these metrics is of practical significance because they are currently in use for regulatory purposes. However in recent work these and other statistical metrics have been evaluated for utility as features for supervised machine learning. In future work we will assess *aer2vec’s* utility as a basis for downstream machine learning also.

## Conclusion

In this paper, we described how *aer2vec* distributed representations of AER data can be used to acquire signal for drug/side-effect associations, outperforming established disproportionality metrics on two pharmacovigilance reference standards. Considering provider designations of suspected causality resulted in further improvements in performance in both *aer2vec* and baseline models. As the baseline disproportionality models concerned are in current use, these results have immediate implications for pharmacovigilance practice, with the potential for broad application of the embeddings that result.

## Acknowledgments

This work was supported by U.S. National Library of Medicine Grant (R01-LM011563), Robust Inference from Observational Data with Distributed Representations of Conceptual Relations.

## Footnotes

i https://github.com/treversec/aer2vec

ii https://zenodol.org/record/3283012

## References

1. Coloma PM, Trifirò G, Patadia V, Sturkenboom M. Postmarketing safety surveillance. Drug Saf. 2013;36(3):183–197.

2. Wysowski DK, Swartz L. Adverse Drug Event Surveillance and Drug Withdrawals in the United States, 1969-2002: The Importance of Reporting Suspected Reactions. Arch Intern Med. 2005 Jun 27;165(12):1363.

3. Research C for DE and. FDA Adverse Events Reporting System (FAERS) - Reports Received and Reports Entered into FAERS by Year[Internet].[cited 2016 Oct 21]. Available from: http://www.fda.gov/Drugs/GuidanceComplianceRegulatoryInformation/Surveillance/AdverseDrugEffects/ucm070434.htm

4. Evans SJW, Waller PC, Davis S. Use of proportional reporting ratios (PRRs) for signal generation from spontaneous adverse drug reaction reports. Pharmacoepidemiol Drug Saf. 2001 Dec 10;10(6):483–6.

5. Heijden VD, M PG, Puijenbroek V, P E, van Buuren S, Hofstede VD, et al. On the assessment of adverse drug reactions from spontaneous reporting systems: the influence of under-reporting on odds ratios. Stat Med. 2002 Jun 19;21(14):2027–44.

6. Bate A, Evans SJW. Quantitative signal detection using spontaneous ADR reporting. Pharmacoepidemiol Drug Saf. 2009 Jun 1;18(6):427–36.

7. Ahmed I, Dalmasso C, Haramburu F, Thiessard F, Broët P, Tubert-Bitter P. False Discovery Rate Estimation for Frequentist Pharmacovigilance Signal Detection Methods. Biometrics. 2009 May 7;66(1):301–9.

8. Bate A, Lindquist M, Edwards IR, Olsson S, Orre R, Lansner A, et al. A Bayesian neural network method for adverse drug reaction signal generation. Eur J Clin Pharmacol. 1998;54(4):315–21.

9. Szarfman A, Machado SG, O’Neill RT. Use of screening algorithms and computer systems to efficiently signal higher-than-expected combinations of drugs and events in the US FDA’s spontaneous reports database. Drug Saf Int J Med Toxicol Drug Exp. 2002;25(6):381–92.

10. Harpaz R, DuMouchel W, LePendu P, Bauer-Mehren A, Ryan P, Shah NH. Performance of Pharmacovigilance Signal Detection Algorithms for the FDA Adverse Event Reporting System. Clin Pharmacol Ther[Internet]. 2013 Jun[cited 2018 Dec 11];93(6). Available from: https://www.ncbi.nlm.nih.gov/pmc/articles/PMC3857139/

11. Harris, Z. (1954). “Distributional structure”. Word. 10(23): 146–162. doi:10.1080/00437956.1954.11659520

12. Cohen T, Widdows D. Empirical distributional semantics: methods and biomedical applications. J Biomed Inform. 2009 Apr;42(2):390–405.

13. Turney PD, Pantel P. From frequency to meaning: Vector space models of semantics. J Artif Intell Res. 2010;37(1):141–188.

14. Mikolov T, Sutskever I, Chen K, Corrado GS, Dean J. Distributed Representations of Words and Phrases and their Compositionality. In: Burges CJC, Bottou L, Welling M, Ghahramani Z, Weinberger KQ, editors. Advances in Neural Information Processing Systems 26[Internet]. Curran Associates, Inc.; 2013[cited 2017 May 31]. p. 3111–3119. Available from: http://papers.nips.cc/paper/5021-distributed-representations-of-words-and-phrases-and-their-compositionality.pdf

15. Mikolov T, Chen K, Corrado G, Dean J. Efficient Estimation of Word Representations in Vector Space. ICLR Workshop 2013; Available from: http://arxiv.org/abs/1301.3781

16. Google Code Archive - Long-term storage for Google Code Project Hosting.[Internet].[cited 2019 Feb 5]. Available from: https://code.google.com/archive/p/word2vec/

17. fastText[Internet].[cited 2019 Feb 5]. Available from: https://fasttext.cc/index.html

18. Rothman KJ, Lanes S, Sacks ST. The reporting odds ratio and its advantages over the proportional reporting ratio. Pharmacoepidemiol Drug Saf. 2004 Aug;13(8):519–23.

19. Tomas Mikolov, Ilya Sutskever, Kai Chen, Greg Corrado, and Jeffrey Dean. 2013b. Distributed representations of words and phrases and their compositionality. In NIPS, pages 3111–3119.

20. Goldberg Y, Levy O. word2vec Explained: deriving Mikolov et al.’s negative-sampling word-embedding method. arXiv preprint arXiv:1402.3722. 2014 Feb 15.

21. Levy O, Goldberg Y, Dagan I. Improving distributional similarity with lessons learned from word embeddings. Transactions of the Association for Computational Linguistics. 2015 Dec;3:211–25.

22. Banda JM, Evans L, Vanguri RS, Tatonetti NP, Ryan PB, Shah NH. A curated and standardized adverse drug event resource to accelerate drug safety research. Sci Data[Internet]. 2016 May 10 [cited 2016 Oct 30];3. Available from: http://www.ncbi.nlm.nih.gov/pmc/articles/PMC4872271/

23. Ryan PB, Schuemie MJ, Welebob E, Duke J, Valentine S, Hartzema AG. Defining a reference set to support methodological research in drug safety. Drug Saf. 2013 Oct;36 Suppl 1:S33–47.

24. Coloma PM, Schuemie MJ, Trifirò G, Gini R, Herings R, Hippisley-Cox J, et al. Combining electronic healthcare databases in Europe to allow for large-scale drug safety monitoring: the EU-ADR Project. Pharmacoepidemiol Drug Saf. 2011 Jan;20(1):1–11.

25. Waller P, Van Puijenbroek EP, Egberts AC, Evans S. The reporting odds ratio versus the proportional reporting ratio:‘deuce’. Pharmacoepidemiology and drug safety. 2004;13(8):525–6

26. Shang N, Xu H, Rindflesch TC, Cohen T. Identifying plausible adverse drug reactions using knowledge extracted from the literature. Journal of Biomedical Informatics. 2014 Dec;52:293–310.

27. Mower J, Subramanian D, Shang N, Cohen T. Classification-by-Analogy: Using Vector Representations of Implicit Relationships to Identify Plausibly Causal Drug/Side-effect Relationships. AMIA Annu Symp Proc. 2017 Feb 10;2016:1940–9.

28. Voss EA, Boyce RD, Ryan PB, van der Lei J, Rijnbeek PR, Schuemie MJ. Accuracy of an automated knowledge base for identifying drug adverse reactions. Journal of biomedical informatics. 2017 Feb 1;66:72–81.

29. McCoy TH, Perlis RH. A tool to utilize adverse effect profiles to identify brain-active medications for repurposing. International Journal of Neuropsychopharmacology. 2015 Feb 1;18(3).

30. Li Y, Ryan PB, Wei Y, Friedman C. A Method to Combine Signals from Spontaneous Reporting Systems and Observational Healthcare Data to Detect Adverse Drug Reactions. Drug Saf. 2015 Oct;38(10):895–908.

31. Malec SA, Wei P, Xu H, Bernstam EV, Myneni S, Cohen T. Literature-Based Discovery of Confounding in Observational Clinical Data. AMIA Annu Symp Proc. 2017 Feb 10;2016:1920–9.

32. Tatonetti NP, Patrick PY, Daneshjou R, Altman RB. Data-driven prediction of drug effects and interactions. Science translational medicine. 2012 Mar 14;4(125):125ra31-.

33. Pariente A, Gregoire F, Fourrier-Reglat A, Haramburu F, Moore N. Impact of safety alerts on measures of disproportionality in spontaneous reporting databases the notoriety bias. Drug safety. 2007 Oct 1;30(10):891–8.

34. Surowiecki J. The wisdom of crowds. Anchor; 2005.

35. Yin Z, Shen Y. On the dimensionality of word embedding. In: Advances in Neural Information Processing Systems 2018 (pp. 887–898).

36. Norén GN, Caster O, Juhlin K, Lindquist M. Zoo or Savannah? Choice of Training Ground for Evidence-Based Pharmacovigilance. Drug Saf. 2014 Sep 1;37(9):655–9.

